# Exploring a natural baseline for large herbivore biomass

**DOI:** 10.1101/2020.02.27.968461

**Authors:** Camilla Fløjgaard, Pil Birkefeldt Pedersen, Christopher J. Sandom, Jens-Christian Svenning, Rasmus Ejrnæs

## Abstract

The massive global losses of large mammals in the Pleistocene have triggered severe ecosystem changes including changed nutrient cycles, fire regimes and climate, shifts in biomes and loss of biodiversity. Large herbivores create and diversify resources and living space for other organisms and thereby play an important role in ecosystem functioning and biodiversity conservation. However, even today large herbivores are regulated, hunted and driven to extinction to a degree where intact large-herbivore communities are largely non-existent. Consequently, natural density and biomass of large-herbivores for restoration of ecosystems are poorly known. To address this knowledge gap, we apply the scaling pattern for consumer-producer relationships and show that the biomass of large herbivores in ecosystems across the world is considerably lower than expected from primary productivity. African ecosystems have the strongest consumer-producer relationship and assuming that African ecosystems approach a natural baseline, we use this relationship to predict large herbivore biomass in Europe as an example. Our findings indicate that restoring large herbivore biomass would entail increasing large herbivore biomass by orders of magnitude in most ecosystems, which potentially changes the perspective on large herbivores in conservation and restoration projects.

## Introduction

The massive losses of large mammals in the late-Quaternary (Faurby & Svenning, 2015) left depauperate large-herbivore communities in most areas (Svenning et al., 2016) and triggered severe ecosystem changes including changed nutrient cycles, climate and fire regimes, shifts in biomes and loss of biodiversity (Doughty et al., 2016; Gill, Williams, Jackson, Lininger, & Robinson, 2009; Malhi et al., 2016). Reinstating large herbivore ecosystem function is becoming increasingly relevant in ecosystem restoration (Fernandez, Navarro, & Pereira, 2017; Perino et al., 2019; Svenning et al., 2016) given their importance in shaping the ecosystem structure, function, composition and diversity (Bakker et al., 2016; Doughty et al., 2016; Estes et al., 2011; Galetti et al., 2018; Malhi et al., 2016). Most species of current ecosystems evolved in prehistoric ecosystems shaped by natural processes including large herbivores (MacFadden, 1997; Sandom, Ejrnæs, Hansen, & Svenning, 2014; Weil, 2005). The ecosystem functions of large herbivores include the diversification of biotic resources and abiotic environment for other species (Doughty, Wolf, & Malhi, 2013; Galetti et al., 2018) thereby contributing to biodiversity conservation.

Today, large mammals are sought out and threatened by regulation, displacement, hunting, and poaching, resulting in wild terrestrial mammals only accounting for 4 % of total mammal biomass today (Bar-On, Phillips, & Milo, 2018; Dirzo et al., 2014). Concurrently, wildlife management literature abounds with claims of overabundance, hyper-abundance and unnaturally high densities of large herbivores. These claims are rarely based on comparison with natural baselines or carrying capacity but rather concluded from anticipated negative effects on valued aspects of ecosystems, such as human life or well-being, the fitness of the large herbivore, the density of species with an economic or aesthetic value, or dysfunctions in the ecosystem (Caughley, 1981). Moreover, this seems a text book case of shifting baseline syndrome (Papworth, Rist, Coad, & Milner-Gulland, 2009; Vera, 2010), as we have forgotten how masses of large mammals used to roam ecosystems across the globe. In order to avoid the danger of managing for preconceived conservation targets and a shifted baseline, we recommend the consideration of natural baselines for the density of large herbivores and to base these on empirical data and experimentation. In this study, we use empirical data and ecological theory to investigate natural baselines for the density of large terrestrial herbivores. We acknowledge that carrying capacity is a complex attribute, but for this preliminary collation and analysis of a global data set, we assume that the collective natural densities of large herbivores are primarily regulated by primary productivity. We recognize that predator-driven top-down regulation can occur, but while predators can limit smaller and intermediate-sized prey, predation tends to cause a community shift towards increased densities of larger bodied herbivores less affected by predation (le Roux et al., 2019). Our assumption follows what has been formulated as a general scaling law in ecology (Cebrian, 2015) describing how energy flows between trophic levels, e.g., predator-prey biomass relationships.

Scaling laws have been demonstrated for many consumer-producer and predator-prey relationships using both biomass and productivity (Cebrian, 2004; Coe, Cumming, & Phillipson, 1976; Gasol, del Giorgio, & Duarte, 1997; Hatton et al., 2015). Hatton et al. (2015) showed that for ecosystems representing near-natural conditions, across trophic levels and ecosystem productivity, there is a strong correlation between producer and consumer biomass typically following a scaling law with an exponent close to ¾. Hatton et al. (2015) accentuate the close relationship between large mammal prey biomass and predator biomass and demonstrate similar exponents for productivity-consumer biomass relationships. Others have also found a close association between rainfall, as an estimate of primary productivity, and large herbivore biomass in Africa with an exponent of c. 1.5-2 (Coe et al., 1976; East, 1984; Fritz & Duncan, 1994).

Given the more intact terrestrial mammal faunas in Africa (Svenning et al., 2016), we expect data from African ecosystems to fit the theoretical scaling between productivity and large herbivore biomass whereas we hypothesize that data from regions heavily influenced by humans reveal lower herbivore biomass and poorer fit. Here, we collate global data on the biomass of large herbivores and combine it with Net Primary Production (NPP) to test the applicability of the general scaling law to large herbivore biomass in terrestrial ecosystems. After exploring the scaling pattern across continents, we use the African scaling pattern to predict large herbivore biomass in Europe as an example and discuss natural baselines of large herbivores for ecosystem restoration.

### Applying the scaling law to large herbivore biomass and primary productivity

We collated published empirical data on large herbivore biomass from several published sources (Baker, Cornelissen, Bhagwat, Vera, & Willis, 2016; Coe et al., 1976; Hatton et al., 2015; Rodriguez et al., 2014) (please see original references therein) and from personal communication (see S1). We included biomass of migrating species and megaherbivores (i.e., herbivores >1000 kg, e.g., elephants, rhinos, and hippos) and areas denoted as wolf-exploited, thereby making some modifications to the sum of large herbivore biomass in Hatton et al. (2015). From Rodriguez et al. (2014) we use all data including ecosystems with livestock grazing. We include these partly pastoral areas because grazing livestock can consume a significant amount of primary production and contribute to the total ungulate biomass. Areas from Coe et al. (1976) were georeferenced from the published map to get coordinates in latitude and longitude otherwise we use the published coordinates. We use data from all years and data from unknown years. While this may cause a temporal mismatch with the yearly range for net primary productivity for some data points (see below), it is unlikely to introduce a systematic bias and we, therefore, prioritize biomass data quantity. In total, we include 317 data points from 169 ecosystems across the five continents of Africa, Asia, Europe, North America and South America (see S2).

As an estimate of primary productivity, we use satellite-derived net primary productivity (Zhao, Heinsch, Nemani, & Running, 2005; Zhao & Running, 2010; Zhao, Running, & Nemani, 2006) (NPP retrieved from http://files.ntsg.umt.edu/data/NTSG_Products/MOD17/GeoTIFF/MOD17A3/GeoTIFF_30arcsec/), which is the average of yearly NPP from 2000 to 2015 in 30 arc seconds (global). We calculate mean and median NPP at different scales using buffers of 1, 5, 10, 50 and 100 km surrounding the given coordinate of the ecosystem.

For ecosystems with the same name, we use the average biomass and NPP (the same ecosystem can have slightly different coordinates depending on the source). Conferring with Hatton et al. (2015), we use ordinary least squares (OLS) to fit the loge-transformed biomass and NPP data. We chose the buffer size with the best fit (highest R^2^, S4) to display global and continental relationships. Building on the assumption that African ecosystems are our most reliable baseline, we construct a model on a subset of African ecosystems with megaherbivores. We use this model to predict large herbivore biomass of Europe. The detailed analysis can be obtained from the available R-script (R Core Team, 2017) (S3).

### Empirical findings of large herbivore biomass

The global data reveals orders of magnitude of differences in large herbivore biomass across ecosystems and between continents with highly irregular scaling patterns (Fig. 1). African and Asian ecosystems have the highest herbivore biomass, whereas North and South America have the lowest. South America also deviates from the general increasing association between productivity and biomass, by having the lowest herbivore biomass at high productivity ecosystems (Fig. 1).

**Figure 1.**
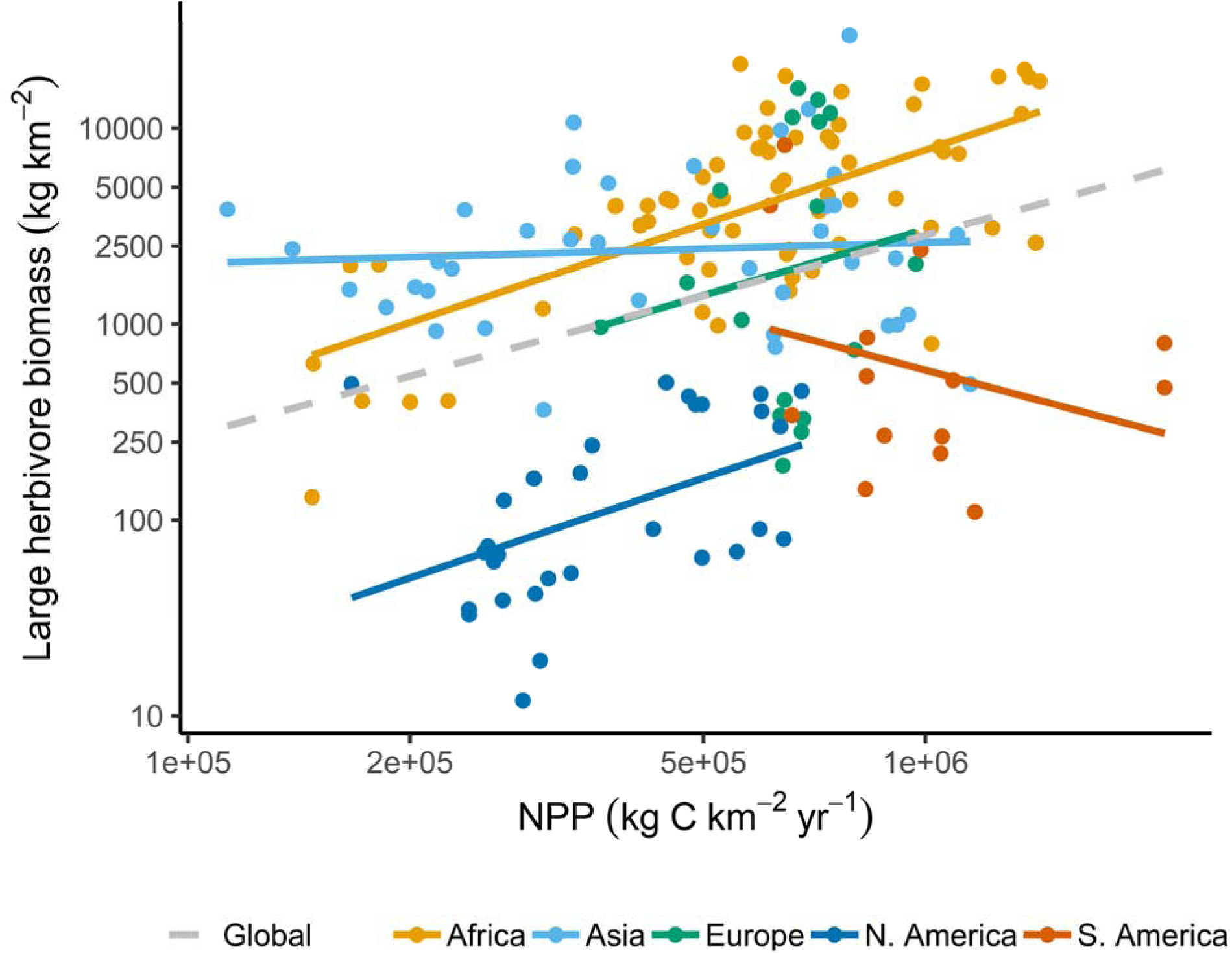
The scaling pattern of large herbivore biomass and NPP. Log-log linear regression lines of large herbivore biomass as a function of primary productivity (mean NPP at buffer scale 100 km). The colors of points and lines show that the relationships and data for the global data or subset by continent.

Ordinary least squares fit to log-transformed data shows a significant, but relatively poor fit for the global data set (R^2^=0.11, n=169, P < 0.001, Table 1). Across African ecosystems (Fig. 1) from some of the most productive ecosystems in the world to the Namib Desert, the large herbivore biomass follows a consistent relationship to ecosystem productivity with the following scaling pattern: Biomass = 0.07*NPP^1.26^ (R^2^=0.43, n=69, P < 0.001, Table 1). Subsetting the data to African ecosystems with megaherbivores increases model fit and yields a scaling pattern of Biomass = 0.01*NPP^1.51^ (R^2^=0.69, n=23, P < 0.001, Table 1). Large herbivore biomass is considerably lower in ecosystems outside Africa and a general scaling pattern is hardly discoverable on other continents (Table 1).

**Table 1.** Summary of ordinary least squares log-log models for large herbivore biomass as a function of primary productivity (NPP). Rows represent geographic regions and the primary productivity (NPP) measure, i.e. median or mean and the scale of the buffer (radius in km) surrounding the ecosystem center coordinate, that yields the best model fit (highest R^2^ in S4). Model significance is indicated by ns, non-significant, * p≤0.05, ** p≤0.01, and *** p≤0.001. The intercept and slope can be used to formulate the scaling law according to Biomass = intercept*NPP^slope^. N denotes the number of data points in the model and current biomass gives the current mean and standard deviation of large herbivore biomass within the regions.

| Continent | NPP calculation and buffer scale | $R^2$ | intercept (log) | intercept | slope | n | Current biomass, mean (kg/km<sup>2</sup>) | Current biomass, sd (kg/km<sup>2</sup>) |
| --- | --- | --- | --- | --- | --- | --- | --- | --- |
| Africa | mean, 100 km | 0.43*** | -2.66 | 0.07 | 1.26 | 69 | 7074 | 6371 |
| Asia | mean, 5 km | -0.02<sup>ns</sup> | 6.94 | 1031.06 | 0.10 | 39 | 3071 | 3694 |
| Europe | mean, 100 km | -0.04<sup>ns</sup> | -2.30 | 0.10 | 1.12 | 17 | 4760 | 5617 |
| NorthAmerica | mean, 50 km | 0.20** | -5.28 | 0.01 | 1.22 | 30 | 208 | 189 |
| SouthAmerica | mean, 100 km | 0.02<sup>ns</sup> | 15.56 | 5718603.32 | -1.00 | 14 | 1371 | 2250 |
| Global | mean, 100 km | 0.11*** | -1.52 | 0.22 | 1.03 | 169 | 3939 | 5249 |
| Africa (megaherbivores) | median, 1 km | 0.69*** | -4.60 | 0.01 | 1.51 | 23 | 6892 | 5418 |

Using the African ecosystems as a baseline, predicted large herbivore biomass of Europe has a mean of 4840 kg/km^2^ and a 5 % quantile range of 875 – 9846 kg/km^2^. This is close to the current mean large herbivore biomass of 4760 kg/km^2^ (SD= 5617, n=17, Table 1). However, the current distribution is strongly bimodal with a peak at high biomass consisting exclusively of rewilding sites and a peak at low herbivore biomass (Fig. 2A). Predicted herbivore biomass with low values coincides with known low-productivity areas, i.e., dry areas of southern Europe, cold regions in northern Europe and in the mountainous regions, and generally high herbivore biomass in central Europe (Fig. 2B).

**Figure 2.**
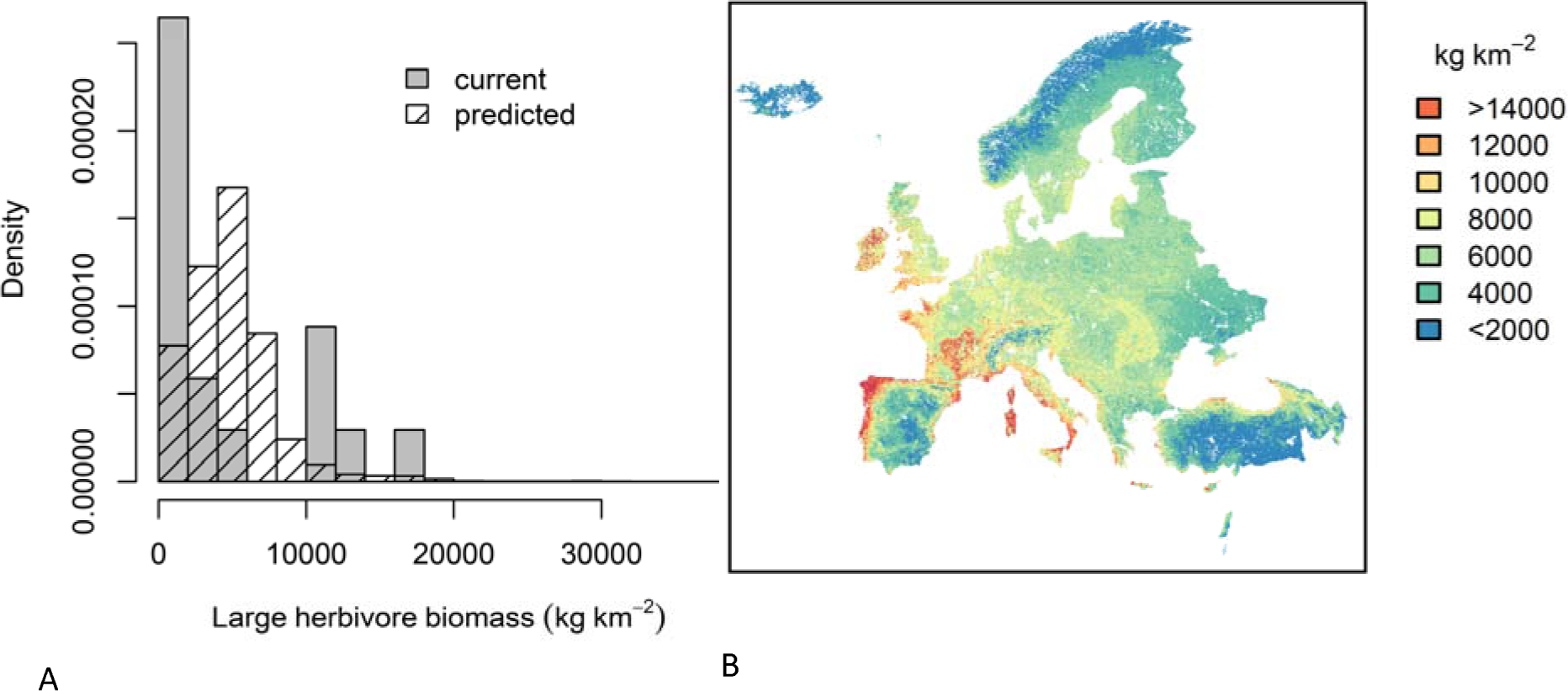
Predicted large herbivore biomass of Europe based on the African scaling pattern as a natural baseline. A. European map of predicted values of large herbivore biomass using the African scaling pattern, i.e., Biomass = 0.01*NPP^1.51^ (R^2^=0.69, n=23, P < 0.001). Biomass increases with warm colors and units are in kg/km^2^. B. Histogram of predicted large herbivore biomass (blue, corresponding to the mapped biomass) and current biomass in Europe (red, n=17).

## Discussion

Our findings of a scaling pattern in African ecosystems is consistent with previous findings, i.e., the general scaling patterns between other trophic levels and across biomes (Hatton et al., 2015) and specifically a scaling pattern between large herbivore biomass and rainfall in African ecosystems (Coe et al., 1976). We find a lower fit in our scaling pattern, which could be attributed to the inclusion of migratory species that consume primary productivity across different ecosystems and the use of satellite-derived NPP. While NPP has the advantage of high-resolution global coverage and broad temporal coverage (mean of 15 years) there can be a discrepancy between NPP and actual available forage, especially in forests.

The latter could be a contributing factor to the – seemingly – low herbivore biomass estimates in the most productive ecosystems in South America, where high productivity areas coincide with rainforests. However, South American fauna suffered massive losses of Pleistocene megafauna (Barnosky, Koch, Feranec, Wing, & Shabel, 2004) and this may also have contributed to relatively low herbivore biomass across all ecosystems in South America. A previous study found that on the rangelands in South America, the introduction of livestock (supplementary feeding and practices such as irrigation and fertilization are rare) increases the large herbivore biomass by an order of magnitude compared to surrounding unmanaged ecosystems (Oesterheld, Sala, & McNaughton, 1992), indicating that the biomass of wild large herbivores is well below carrying capacity.

Although North America appears to follow a scaling law with a significant positive relationship between productivity and large herbivore biomass (Table 1), the biomass is consistently low across the continent. This could indicate that the extensive Late Pleistocene megafauna extinctions (Barnosky et al., 2004; Gill et al., 2009) were associated with both low- and high productive ecosystems causing a general decrease in biomass. Moreover, extant species may persist at low densities, e.g., a simulation study has shown that bison in Yellowstone NP has not yet reached the estimated mean food-limited carrying capacity of the area (Plumb, White, Coughenour, & Wallen, 2009).

There seems to be no scaling pattern between productivity and large herbivore biomass in the data from Asian ecosystems. Extinction of megafauna and suppressed fire regimes, causing a decrease in palatable forage in woodlands, are some of the suggested explanations for seemingly low large herbivore biomass in some areas (Karanth & Sunquist, 1992). Poaching may also notably reduce large herbivore biomass (Ripple et al., 2016), e.g., contrasting biomass of 1450 kg/km^2^ (Srikosamatara, 1993) and 14,744 kg/km^2^ (Karanth & Sunquist, 1992) in two dry tropical forest ecosystems was seemingly due to intensive poaching activity and lack of wildlife management at the first site.

There is considerable variation in large herbivore biomass in Europe, ranging from a minimum of 190 kg/km^2^ to 16,000 kg/km^2^ with a notable bimodal distribution (Fig. 2A). The highest figures are derived from rewilding areas, such as Rewilding Europe’s Lika Plains, FreeNature’s areas at Geldersee Port and the Natural History Museum Aarhus’ Mols Laboratory (Table 2), that all practice ‘natural grazing’, i.e., cattle (*Bos taurus*) and horse (*Equus caballus*) have been introduced to (often fenced) nature reserves, where their densities are allowed to increase and are only regulated to avoid winter starvation (Jepson, Schepers, & Helmer, 2018). In practice, this approaches populations regulated by primary productivity.

**Table 2.**
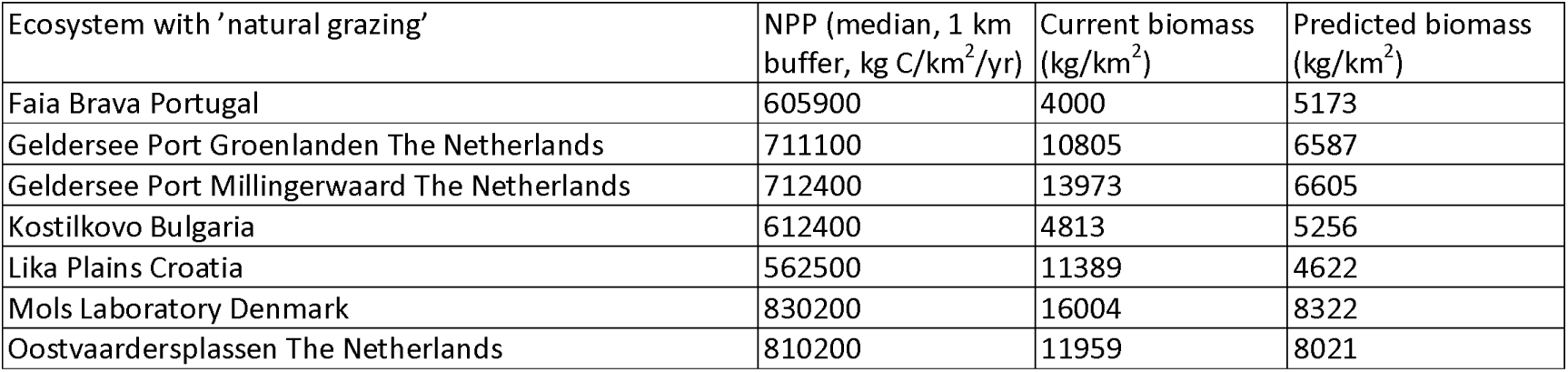
Current and predicted large herbivore biomass for European ecosystems and nature areas known to practice ‘natural grazing’. Rows represent geographic regions, their primary productivity (NPP), and current and predicted large herbivore biomass based on the African scaling pattern, i.e., Biomass = 0.01*NPP^1.51^ (R^2^=0.69, n=23, P < 0.001).

Based on the African scaling pattern, large herbivore biomass of up to 9846 kg/km^2^ (95% quantile) is predicted across European ecosystems (Fig. 2A). However, those few nature areas that practice ‘natural grazing’ reach large herbivore biomasses within or above this prediction (Fig. 2A, Table 2), indicating that the prediction could be somewhat conservative. Notably, the Oostvaardersplassen (OVP), a 5600 ha Dutch rewilding site allowing unregulated population growth of fenced cattle, horse and red deer (*Cervus elaphus*), reached higher large herbivore biomass with fluctuating large herbivore densities of up to c. 12,000 kg/km^2^ (in effect even higher considering that a large part of the reserve is permanently flooded and unsuitable for these large herbivores). Also, palaeoecological data yields higher estimates than the prediction, e.g., the mammoth steppe is estimated to have had a megafauna biomass of 10,500 kg/km^2^ (Zimov, Zimov, Tikhonov, & Chapin, 2012) and in Holocene England 2.5 large herbivores/ha is estimated to have roamed (Smith, Whitehouse, Bunting, & Chapman, 2009), i.e., 25,000 kg/km^2^ if the large herbivores weighed 100 kg as an example.

European ecosystems are obviously lacking key large herbivores and megaherbivores (Bunzel-Drüke, Drüke, & Vierhaus, 2001) – not just extinct species. Extant species, such as the European bison (*Bison bonasus*), today only occupy a fraction of their previous distribution, with repercussions for vegetation and biodiversity. Sandom et al. (2014) showed that 55 % of Last Interglacial sites had a high number of dung-indicator beetles and beetles corresponding to widespread wood pastures. Concordantly, there is an increasing focus on reintroducing missing species to nature areas (Pedersen, Ejrnæs, Sandel, & Svenning, 2019), however, given the scaling pattern, there is grounds to also focus on restoring ecosystem functions by allowing natural densities of large herbivores to develop.

Reestablishing large herbivore ecosystem function naturally leads to discussions of the role of predators and it is argued that predation will reduce the densities of large herbivores and that reintroduction of predators may solve problems associated with higher grazing and browsing pressure (Bull, Ejrnaes, Macdonald, Svenning, & Sandom, 2018; Nilsen et al., 2007; Sandom, Bull, Canney, & Macdonald, 2012). Predation can play an important role in the spatial distribution and behavior of herbivores (Laundré, Hernández, & Ripple, 2010; Ripple & Beschta, 2004) and cause a notable shift in the biomass distribution of herbivores towards large bodied species (Hopcraft, Anderson, Pérez-Vila, Mayemba, & Olff, 2012; Ripple, Painter, Beschta, & Gates, 2011), but evidence that predators generally and substantially regulate total herbivore densities is at best equivocal and varies across ecosystems (Hopcraft, Olff, & Sinclair, 2010; Skogland, 1991). In other words, there are grounds to focus on ecosystem restoration from multiple aspects of large herbivores, i.e., introducing relevant species, notably large- and megaherbivores, increasing biomass and reestablishing predation (Svenning, 2019).

### Perspectives for establishing a natural baseline

Outside African ecosystems, large herbivore biomass does not follow the general scaling pattern with primary productivity and is generally much lower than expected from primary productivity. Using African data as a baseline much higher biomass could be expected in most nature areas in Europe, except for rewilding areas and palaeoecological estimates that tend to be in the high range of or even exceed the predictions. We can easily explain densities below predictions by population regulation and past defaunation, but densities exceeding predictions point to unresolved issues. Most rewilding areas in Europe are relatively young, and some but not all are without predators, and it is possible that herbivore populations will settle at a lower biomass in the end. The effects of interactions between large predators (Hopcraft et al., 2010; Skogland, 1991), other species of large herbivores and megaherbivores in more intact mammal assemblages (Tanentzap & Smith, 2018), feedback from vegetation (Valeix et al., 2011) and climate change (Olofsson & Post, 2018) on the overall large herbivore biomass is also hard to predict. Another explanation for the discrepancy between predicted and observed biomass of rewilding areas, may be that the African ecosystems are not as near-natural as expected, i.e., also here, large herbivore populations are reduced by e.g. poaching, culling or immunocontraception of elephants (Fayrer-Hosken, Grobler, Van Altena, Bertschinger, & Kirkpatrick, 2000). It is hard to exclude the possibility that experimental European rewilding sites with ‘natural grazing’ allowing population growth are closer to the natural baseline of large herbivore biomass.

Late Pleistocene extinctions and the current global defaunation have severe impact on biodiversity (Bakker et al., 2016; Sandom et al., 2014). Moreover, it has contributed to a shifting baseline in conservation, leading to wildlife management targeting subjective human perceptions of ecosystems (Papworth et al., 2009; Vera, 2010). Starting from qualified guesses based on empirical data and scaling law, we propose that further progress in establishing baselines requires large-scale field experiments where diverse assemblages of large herbivores are allowed to respond to primary productivity and seasonality without proactive regulation. We hope this study can be a starting point of research in large herbivore baselines. For example, a network of experimental rewilding areas (Jepson, 2016) seeking to restore large herbivore and predator assemblages to natural densities.

## Supporting information

Data from personal communication

Full data set

R script

Model results

## Acknowledgements

We are grateful to donations to CF and RE from Klelund Deer Park and Aage V Jensens Fonde (DNI-project). JCS considers this work a contribution to his Carlsberg Foundation Semper Ardens project MegaPast2Future (grant CF16-0005) and to his VILLUM Investigator project “Biodiversity Dynamics in a Changing World” funded by VILLUM FONDEN (grant 16549). PBP considers this work a contribution to her Carlsberg Foundation Internationalization Fellowship.

## Supplementary Information

**Appendix S1 – Data from personal communication**

**Appendix S2 – Full data set of large herbivore biomass for analysis**

**Appendix S3 – R script for data analysis**

**Appendix S4 – Full table of regression results of large herbivore biomass and NPP**

## References

Baker, A. G., Cornelissen, P., Bhagwat, S. A., Vera, F. W., & Willis, K. J. (2016). Quantification of population sizes of large herbivores and their long-term functional role in ecosystems using dung fungal spores. Methods in Ecology and Evolution.

Bakker, E. S., Gill, J. L., Johnson, C. N., Vera, F. W., Sandom, C. J., Asner, G. P., & Svenning, J.-C. (2016). Combining paleo-data and modern exclosure experiments to assess the impact of megafauna extinctions on woody vegetation. Proceedings of the National Academy of Sciences, 113(4), 847–855.

Bar-On, Y. M., Phillips, R., & Milo, R. (2018). The biomass distribution on Earth. Proc Natl Acad Sci U S A, 115(25), 6506–6511. doi:10.1073/pnas.1711842115

Barnosky, A. D., Koch, P. L., Feranec, R. S., Wing, S. L., & Shabel, A. B. (2004). Assessing the Causes of Late Pleistocene Extinctions on the Continents. Science, 306(5693), 70–75. doi:10.1126/science.1101476

Bull, J. W., Ejrnaes, R., Macdonald, D. W., Svenning, J.-C., & Sandom, C. J. (2018). Fences can support restoration in human-dominated ecosystems when rewilding with large predators. Restoration Ecology, 0(0). doi:10.1111/rec.12830

Bunzel-Drüke, M., Drüke, J., & Vierhaus, H. (2001). Der Einfluss von Großherbivoren auf die Naturlandschaft Mitteleuropas. Verfügbar in: http://www.abunaturschutz.de/veroeffe.

Caughley, G. (1981). Overpopulation. In S. H. a. D. H. PA Jewell (Ed.), Problems in Management of Locally Abundant Wild Mammals (pp. 7–19): Academic Press: New York.

Cebrian, J. (2004). Role of first-order consumers in ecosystem carbon flow. Ecology Letters, 7(3), 232–240. doi:10.1111/j.1461-0248.2004.00574.x

Cebrian, J. (2015). Energy flows in ecosystems. Science, 349(6252), 1053–1054.

Coe, M. J., Cumming, D. H., & Phillipson, J. (1976). Biomass and Production of Large African Herbivores in Relation to Rainfall and Primary Production. Oecologia, 22(4), 341–354.

Dirzo, R., Young, H. S., Galetti, M., Ceballos, G., Isaac, N. J., & Collen, B. (2014). Defaunation in the Anthropocene. Science, 345(6195), 401–406.

Doughty, C. E., Roman, J., Faurby, S., Wolf, A., Haque, A., Bakker, E. S., … Svenning, J.-C. (2016). Global nutrient transport in a world of giants. Proceedings of the National Academy of Sciences, 113(4), 868–873. doi:10.1073/pnas.1502549112

Doughty, C. E., Wolf, A., & Malhi, Y. (2013). The legacy of the Pleistocene megafauna extinctions on nutrient availability in Amazonia. Nature Geoscience, 6(9), 761–764. doi:10.1038/Ngeo1895

East, R. (1984). Rainfall, soil nutrient status and biomass of large African savanna mammals. African Journal of Ecology, 22(4), 245–270. doi:doi:10.1111/j.1365-2028.1984.tb00700.x

Estes, J. A., Terborgh, J., Brashares, J. S., Power, M. E., Berger, J., Bond, W. J., … Wardle, D. A. (2011). Trophic Downgrading of Planet Earth. Science, 333(6040), 301–306. doi:10.1126/science.1205106

Faurby, S., & Svenning, J. C. (2015). Historic and prehistoric human-driven extinctions have reshaped global mammal diversity patterns. Diversity and Distributions, 21(10), 1155–1166.

Fayrer-Hosken, R. A., Grobler, D., Van Altena, J. J., Bertschinger, H. J., & Kirkpatrick, J. F. (2000). Immunocontraception of African elephants. Nature, 407(6801), 149–149. doi:10.1038/35025136

Fernandez, N., Navarro, L. M., & Pereira, H. M. (2017). Rewilding: A Call for Boosting Ecological Complexity in Conservation. Conservation Letters, 10(3), 276–278. doi:10.1111/conl.12374

Fritz, H., & Duncan, P. (1994). On the carrying capacity for large ungulates of African savanna ecosystems. Proceedings of the Royal Society of London. Series B: Biological Sciences, 256(1345), 77–82. doi:doi:10.1098/rspb.1994.0052

Galetti, M., Moleón, M., Jordano, P., Pires, M. M., Guimarães Jr., P. R., Pape, T., … Svenning, J.-C. (2018). Ecological and evolutionary legacy of megafauna extinctions. Biological Reviews, 93(2), 845–862. doi:10.1111/brv.12374

Gasol, J. M., del Giorgio, P. A., & Duarte, C. M. (1997). Biomass distribution in marine planktonic communities. Limnology and Oceanography, 42(6), 1353–1363. doi:10.4319/lo.1997.42.6.1353

Gill, J. L., Williams, J. W., Jackson, S. T., Lininger, K. B., & Robinson, G. S. (2009). Pleistocene Megafaunal Collapse, Novel Plant Communities, and Enhanced Fire Regimes in North America. Science, 326(5956), 1100–1103. doi:10.1126/science.1179504

Hatton, I. A., McCann, K. S., Fryxell, J. M., Davies, T. J., Smerlak, M., Sinclair, A. R. E., & Loreau, M. (2015). The predator-prey power law: Biomass scaling across terrestrial and aquatic biomes. Science, 349(6252). doi:10.1126/science.aac6284

Hopcraft, J. G., Olff, H., & Sinclair, A. R. (2010). Herbivores, resources and risks: alternating regulation along primary environmental gradients in savannas. Trends in Ecology & Evolution, 25(2), 119–128. doi:10.1016/j.tree.2009.08.001

Hopcraft, J. G. C., Anderson, T. M., Pérez-Vila, S., Mayemba, E., & Olff, H. (2012). Body size and the division of niche space: food and predation differentially shape the distribution of Serengeti grazers. Journal of Animal Ecology, 81(1), 201–213. doi:10.1111/j.1365-2656.2011.01885.x

Jepson, P. (2016). A rewilding agenda for Europe: creating a network of experimental reserves. Ecography, 39(2), 117–124. doi:10.1111/ecog.01602

Jepson, P., Schepers, F., & Helmer, W. (2018). Governing with nature: a European perspective on putting rewilding principles into practice. Philosophical Transactions of the Royal Society B: Biological Sciences, 373(1761), 20170434. doi:10.1098/rstb.2017.0434

Karanth, K. U., & Sunquist, M. E. (1992). Population Structure, Density and Biomass of Large Herbivores in the Tropical Forests of Nagarahole, India. Journal of Tropical Ecology, 8(1), 21–35.

Laundré, J. W., Hernández, L., & Ripple, W. J. (2010). The landscape of fear: ecological implications of being afraid. Open Ecology Journal, 3, 1–7.

le Roux, E., Marneweck, D. G., Clinning, G., Druce, D. J., Kerley, G. I. H., & Cromsigt, J. P. G. M. (2019). Top– down limits on prey populations may be more severe in larger prey species, despite having fewer predators. Ecography, 42(6), 1115–1123. doi:10.1111/ecog.03791

MacFadden, B. J. (1997). Origin and evolution of the grazing guild in New World terrestrial mammals. Trends in Ecology & Evolution, 12(5), 182–187.

Malhi, Y., Doughty, C. E., Galetti, M., Smith, F. A., Svenning, J.-C., & Terborgh, J. W. (2016). Megafauna and ecosystem function from the Pleistocene to the Anthropocene. Proceedings of the National Academy of Sciences, 113(4), 838–846. doi:10.1073/pnas.1502540113

Nilsen, E. B., Milner-Gulland, E. J., Schofield, L., Mysterud, A., Stenseth, N. C., & Coulson, T. (2007). Wolf reintroduction to Scotland: public attitudes and consequences for red deer management. Proceedings of the Royal Society B: Biological Sciences, 274(1612), 995–1003. doi:10.1098/rspb.2006.0369

Oesterheld, M., Sala, O. E., & McNaughton, S. J. (1992). Effect of animal husbandry on herbivore-carrying capacity at a regional scale. Nature, 356(6366), 234–236. doi:10.1038/356234a0

Olofsson, J., & Post, E. (2018). Effects of large herbivores on tundra vegetation in a changing climate, and implications for rewilding. Philosophical Transactions of the Royal Society B: Biological Sciences, 373(1761), 20170437.

Papworth, S. K., Rist, J., Coad, L., & Milner-Gulland, E. J. (2009). Evidence for shifting baseline syndrome in conservation. Conservation Letters, 2(2), 93–100. doi:10.1111/j.1755-263X.2009.00049.x

Pedersen, P. B. M., Ejrnæs, R., Sandel, B., & Svenning, J.-C. (2019). Trophic Rewilding Advancement in Anthropogenically Impacted Landscapes (TRAAIL): A framework to link conventional conservation management and rewilding. Ambio, 1–14.

Perino, A., Pereira, H. M., Navarro, L. M., Fernández, N., Bullock, J. M., Ceauşu, S., … Wheeler, H. C. (2019). Rewilding complex ecosystems. Science, 364(6438), eaav5570. doi:10.1126/science.aav5570

Plumb, G. E., White, P. J., Coughenour, M. B., & Wallen, R. L. (2009). Carrying capacity, migration, and dispersal in Yellowstone bison. Biological Conservation, 142(11), 2377–2387. doi:10.1016/j.biocon.2009.05.019

R Core Team. (2017). R: A language and environment for statistical computing. Vienna, Austria: R Foundation for Statistical Computing. Retrieved from http://www.R-project.org

Ripple, W. J., Abernethy, K., Betts, M. G., Chapron, G., Dirzo, R., Galetti, M., … Machovina, B. (2016). Bushmeat hunting and extinction risk to the world’s mammals. Royal Society open science, 3(10), 160498.

Ripple, W. J., & Beschta, R. L. (2004). Wolves and the ecology of fear: can predation risk structure ecosystems? Bioscience, 54(8), 755–766.

Ripple, W. J., Painter, L. E., Beschta, R. L., & Gates, C. C. (2011). Wolves, elk, bison, and secondary trophic cascades in Yellowstone National Park. The Open Ecology Journal, 3(1).

Rodriguez, J., Blain, H. A., Mateos, A., Martin-Gonzalez, J. A., Cuenca-Bescos, G., & Rodriguez-Gomez, G. (2014). Ungulate carrying capacity in Pleistocene Mediterranean ecosystems: Evidence from the Atapuerca sites. Palaeogeography Palaeoclimatology Palaeoecology, 393, 122–134. doi:10.1016/j.palaeo.2013.11.011

Sandom, C., Bull, J., Canney, S., & Macdonald, D. W. (2012). Exploring the Value of Wolves (Canis lupus) in Landscape-Scale Fenced Reserves for Ecological Restoration in the Scottish Highlands. In M. J. Somers & M. Hayward (Eds.), Fencing for Conservation: Restriction of Evolutionary Potential or a Riposte to Threatening Processes? (pp. 245–276). New York, NY: Springer New York.

Sandom, C. J., Ejrnæs, R., Hansen, M. D., & Svenning, J. C. (2014). High herbivore density associated with vegetation diversity in interglacial ecosystems. Proc Natl Acad Sci U S A, 111(11), 4162–4167. doi:10.1073/pnas.1311014111

Skogland, T. (1991). What Are the Effects of Predators on Large Ungulate Populations? Oikos, 61(3), 401–411. doi:10.2307/3545248

Smith, D., Whitehouse, N., Bunting, J. M., & Chapman, H. (2009). Can we characterise ‘openness’ in the Holocene palaeoenvironmental record? Modern analogue studies of insect faunas and pollen spectra from Dunham Massey deer park and Epping Forest, England. The Holocene, 20(2), 215–229. doi:10.1177/0959683609350392

Srikosamatara, S. (1993). Density and biomass of large herbivores and other mammals in a dry tropical forest, western Thailand. Journal of Tropical Ecology, 9(1), 33–43. doi:10.1017/S026646740000691X

Svenning, J.-C., Munk, M. & Schweiger, A. (2019). Trophic rewilding – ecological restoration of top-down trophic interactions to promote self-regulating biodiverse ecosystems. In N. P. Johan T. du Toit, & Sarah M. Durant (Ed.), Rewilding Ecological Reviews (pp. 73–98): Cambridge University Press.

Svenning, J.-C., Pedersen, P. B. M., Donlan, C. J., Ejrnæs, R., Faurby, S., Galetti, M., … Vera, F. W. M. (2016). Science for a wilder Anthropocene: Synthesis and future directions for trophic rewilding research. Proceedings of the National Academy of Sciences, 113(4), 898–906. doi:10.1073/pnas.1502556112

Tanentzap, A. J., & Smith, B. R. (2018). Unintentional rewilding: lessons for trophic rewilding from other forms of species introductions. Philosophical Transactions of the Royal Society B: Biological Sciences, 373(1761), 20170445.

Valeix, M., Fritz, H., Sabatier, R., Murindagomo, F., Cumming, D., & Duncan, P. (2011). Elephant-induced structural changes in the vegetation and habitat selection by large herbivores in an African savanna. Biological Conservation, 144(2), 902–912. doi: https://doi.org/10.1016/j.biocon.2010.10.029

Vera, F. (2010). The shifting baseline syndrome in restoration ecology. In M. Hall (Ed.), Restoration and history: the search for a usable environmental past. Edited by M. Hall. Routledge, New York (pp. 98–110).

Weil, A. (2005). Mammalian palaeobiology: Living large in the Cretaceous. Nature, 433(7022), 116.

Zhao, M., Heinsch, F. A., Nemani, R. R., & Running, S. W. (2005). Improvements of the MODIS terrestrial gross and net primary production global data set. Remote Sensing of Environment, 95(2), 164–176.

Zhao, M., & Running, S. W. (2010). Drought-Induced Reduction in Global Terrestrial Net Primary Production from 2000 Through 2009. Science, 329(5994), 940–943. doi:10.1126/science.1192666

Zhao, M., Running, S. W., & Nemani, R. R. (2006). Sensitivity of Moderate Resolution Imaging Spectroradiometer (MODIS) terrestrial primary production to the accuracy of meteorological reanalyses. Journal of Geophysical Research: Biogeosciences, 11(G1).

Zimov, S. A., Zimov, N. S., Tikhonov, A. N., & Chapin, F. S. (2012). Mammoth steppe: a high-productivity phenomenon. Quaternary Science Reviews, 57, 26–45. doi:10.1016/j.quascirev.2012.10.005

