## Supplementary material for "Exploring a natural baseline for large herbivore biomass": Model results

**Supporting Information S4 – Full table of regression results of large herbivore biomass and NPP**

**Table S1**. **All ordinary least squares log-log models for large herbivore biomass as a function of primary productivity (NPP).**

Rows represent geographic regions and the primary productivity (NPP) measure, i.e. median or mean and the scale of the buffer (radius in km) surrounding the ecosystem center coordinate. Model fit is given by R^2^ and significance is indicated by ns, non significant, * p≤0.05, ** p≤0.01, and *** p≤0.001. The intercept and slope can be used to formulate the scaling law according to logBiomass = intercept logNPP^slope^. N denotes the number of data points in the model.

| Continent | NPP | R^2^ | intercept (log-transformed) | intercept | slope | N |
| --- | --- | --- | --- | --- | --- | --- |
| global | mean, 1 km | 0.06*** | 2.00 | 7.39 | 0.62 | 169 |
| global | mean, 5 km | 0.08*** | 0.96 | 2.60 | 0.74 | 169 |
| global | mean, 10 km | 0.08*** | 0.69 | 1.99 | 0.77 | 169 |
| global | mean, 50 km | 0.10*** | -0.86 | 0.42 | 0.95 | 169 |
| global | mean, 100 km | 0.11*** | -1.52 | 0.22 | 1.03 | 169 |
| global | median, 1 km | 0.06** | 2.23 | 9.27 | 0.60 | 169 |
| global | median, 5 km | 0.07*** | 1.14 | 3.14 | 0.72 | 169 |
| global | median, 10 km | 0.07*** | 1.01 | 2.76 | 0.73 | 169 |
| global | median, 50 km | 0.09*** | -0.30 | 0.74 | 0.89 | 169 |
| global | median, 100 km | 0.10*** | -0.85 | 0.43 | 0.95 | 169 |
| Africa | mean, 1 km | 0.33*** | 0.05 | 1.05 | 0.96 | 69 |
| Africa | mean, 5 km | 0.37*** | -1.30 | 0.27 | 1.12 | 69 |
| Africa | mean, 10 km | 0.39*** | -1.53 | 0.22 | 1.14 | 69 |
| Africa | mean, 50 km | 0.42*** | -2.16 | 0.12 | 1.21 | 69 |
| Africa | mean, 100 km | 0.43*** | -2.66 | 0.07 | 1.26 | 69 |
| Africa | median, 1 km | 0.33*** | 0.25 | 1.28 | 0.94 | 69 |
| Africa | median, 5 km | 0.37*** | -0.91 | 0.40 | 1.07 | 69 |
| Africa | median, 10 km | 0.38*** | -1.05 | 0.35 | 1.09 | 69 |
| Africa | median, 50 km | 0.41*** | -1.56 | 0.21 | 1.14 | 69 |
| Africa | median, 100 km | 0.42*** | -2.14 | 0.12 | 1.20 | 69 |
| Asia | mean, 1 km | -0.02^ns^ | 7.30 | 1480.02 | 0.06 | 39 |
| Asia | mean, 5 km | -0.02^ns^ | 6.94 | 1031.06 | 0.10 | 39 |
| Asia | mean, 10 km | -0.02^ns^ | 6.97 | 1065.77 | 0.10 | 39 |
| Asia | mean, 50 km | -0.02^ns^ | 6.79 | 884.89 | 0.12 | 39 |
| Asia | mean, 100 km | -0.02^ns^ | 6.89 | 978.01 | 0.11 | 39 |
| Asia | median, 1 km | -0.03^ns^ | 7.52 | 1848.42 | 0.03 | 39 |
| Asia | median, 5 km | -0.02^ns^ | 7.21 | 1356.08 | 0.07 | 39 |
| Asia | median, 10 km | -0.03^ns^ | 7.49 | 1796.17 | 0.03 | 39 |
| Asia | median, 50 km | -0.02^ns^ | 7.09 | 1194.60 | 0.08 | 39 |
| Asia | median, 100 km | -0.03^ns^ | 7.41 | 1655.35 | 0.04 | 39 |
| Europe | mean, 1 km | -0.07^ns^ | 9.28 | 10719.83 | -0.20 | 17 |
| Europe | mean, 5 km | -0.05^ns^ | 14.89 | 2927730.23 | -0.84 | 17 |
| Europe | mean, 10 km | -0.06^ns^ | 12.91 | 405090.05 | -0.61 | 17 |
| Europe | mean, 50 km | -0.07^ns^ | 5.35 | 209.78 | 0.25 | 17 |
| Europe | mean, 100 km | -0.04^ns^ | -2.30 | 0.10 | 1.12 | 17 |
| Europe | median, 1 km | -0.07^ns^ | 6.82 | 912.15 | 0.08 | 17 |
| Europe | median, 5 km | -0.07^ns^ | 9.75 | 17150.35 | -0.25 | 17 |
| Europe | median, 10 km | -0.06^ns^ | 11.93 | 151058.27 | -0.50 | 17 |
| Europe | median, 50 km | -0.07^ns^ | 5.57 | 262.81 | 0.22 | 17 |
| Europe | median, 100 km | -0.05^ns^ | 0.00 | 1.00 | 0.86 | 17 |
| NorthAmerica | mean, 1 km | 0.02^ns^ | 1.87 | 6.51 | 0.35 | 30 |
| NorthAmerica | mean, 5 km | 0.09^ns^ | -0.55 | 0.58 | 0.64 | 30 |
| NorthAmerica | mean, 10 km | 0.11* | -1.26 | 0.28 | 0.73 | 30 |
| NorthAmerica | mean, 50 km | 0.20** | -5.28 | 0.01 | 1.22 | 30 |
| NorthAmerica | mean, 100 km | 0.18* | -5.81 | 0.00 | 1.28 | 30 |
| NorthAmerica | median, 1 km | 0.05^ns^ | 1.29 | 3.64 | 0.42 | 30 |
| NorthAmerica | median, 5 km | 0.09^ns^ | -0.12 | 0.88 | 0.59 | 30 |
| NorthAmerica | median, 10 km | 0.09^ns^ | -0.27 | 0.76 | 0.61 | 30 |
| NorthAmerica | median, 50 km | 0.20** | -5.61 | 0.00 | 1.25 | 30 |
| NorthAmerica | median, 100 km | 0.19* | -6.14 | 0.00 | 1.31 | 30 |
| SouthAmerica | mean, 1 km | -0.04^ns^ | 12.21 | 201450.99 | -0.63 | 14 |
| SouthAmerica | mean, 5 km | -0.03^ns^ | 12.59 | 294045.67 | -0.67 | 14 |
| SouthAmerica | mean, 10 km | -0.02^ns^ | 12.82 | 371073.68 | -0.70 | 14 |
| SouthAmerica | mean, 50 km | 0.00^ns^ | 14.29 | 1599695.68 | -0.86 | 14 |
| SouthAmerica | mean, 100 km | 0.02^ns^ | 15.56 | 5718603.32 | -1.00 | 14 |
| SouthAmerica | median, 1 km | -0.04^ns^ | 12.00 | 163534.56 | -0.61 | 14 |
| SouthAmerica | median, 5 km | -0.03^ns^ | 12.39 | 239264.97 | -0.65 | 14 |
| SouthAmerica | median, 10 km | -0.03^ns^ | 12.35 | 231594.08 | -0.65 | 14 |
| SouthAmerica | median, 50 km | -0.02^ns^ | 13.61 | 811918.58 | -0.78 | 14 |
| SouthAmerica | median, 100 km | -0.03^ns^ | 13.17 | 525722.90 | -0.74 | 14 |
| Africa (megaherbivores) | mean, 1 km | 0.69*** | -4.81 | 0.01 | 1.54 | 23 |
| Africa (megaherbivores) | mean, 5 km | 0.58*** | -4.79 | 0.01 | 1.54 | 23 |
| Africa (megaherbivores) | mean, 10 km | 0.58*** | -4.51 | 0.01 | 1.51 | 23 |
| Africa (megaherbivores) | mean, 50 km | 0.50*** | -3.94 | 0.02 | 1.43 | 23 |
| Africa (megaherbivores) | mean, 100 km | 0.49*** | -4.26 | 0.01 | 1.47 | 23 |
| Africa (megaherbivores) | median, 1 km | 0.69*** | -4.60 | 0.01 | 1.51 | 23 |
| Africa (megaherbivores) | median, 5 km | 0.57*** | -4.43 | 0.01 | 1.50 | 23 |
| Africa (megaherbivores) | median, 10 km | 0.57*** | -3.89 | 0.02 | 1.44 | 23 |
| Africa (megaherbivores) | median, 50 km | 0.42*** | -2.61 | 0.07 | 1.28 | 23 |
| Africa (megaherbivores) | median, 100 km | 0.47*** | -3.80 | 0.02 | 1.42 | 23 |
